# Temporal Dynamics of the Neural Representation of Social Relationships

**DOI:** 10.1101/856484

**Authors:** Sarah L. Dziura, James C. Thompson

## Abstract

Humans can rapidly encode information from faces to support social judgments and facilitate interactions with others. We can also recall complex knowledge about those individuals, such as their social relationships with others, but the timecourse of this process has not been examined in detail. This study addressed the temporal dynamics of emerging visual and social relationship information using electroencephalography (EEG) and representational similarity analysis (RSA). Participants became familiar with a 10-person social network, and were then shown faces of that network’s members while EEG was recorded. To examine the temporal dynamics of the cognitive processes related to face perception, we compared the similarity structure of neural pattern responses to models of visual processing, face shape similarity, person identity, and social connectedness. We found that all types of information are associated with neural patterns after a face is seen. Visual and identity models became significant early after image onset, but only the identity model stayed associated with neural patterns until 400 ms. Models representing social connections were also present beginning around 200 ms, even in the absence of an explicit task to think about the friendships among the network members. A partial correlation showed that visual and social information contribute uniquely to person perception, although differences were found between models of social connection. This study highlights the speed and salience of social information relating to group dynamics that are present in the brain during person perception.

**Significance Statement:** We live our lives in social groups where complex relationships form among and around us. It is likely that some of the information about social relationships that we observe is integral during person perception, to better help us interact in differing situations with a variety of people. However, when exactly this information becomes relevant has been unclear. In this study, we present evidence that information reflecting observed relationships among a social network is spontaneously represented in whole-brain patterns shortly following visual perception, and is uniquely present around 400 ms. These results are consistent with neuroimaging studies showing spontaneous spatial representation of social network characteristics, and contribute novel insights into the timing of these neural processes.

---

In a socially interconnected world, the ability to perceive and understand complex information about other people is critical. Humans rely heavily on faces to provide social information, and we can make rapid judgments about the age, sex, trustworthiness, and identity of another within a few hundred milliseconds of viewing a face (Dobs et al., 2019; Todorov et al., 2009; Young and Burton, 2018). Recently, it has been shown that information about the social connections and network positions of those we know is represented in a distributed set of brain regions, including inferior parietal, superior temporal, and medial prefrontal cortices (Parkinson et al., 2017; Morelli et al., 2018). However, this work has so far not revealed how such representations unfold over time. Rapid encoding of information about the social connections of others could be valuable during social interactions, as it can help us know who to trust, who might have access to resources we need, or who might be a good source of social support (Feldman-Hall, 2017; Sutcliffe et al., 2012; Zerubavel et al., 2015).

Cognitive (Barry et al., 1998; Bruce and Young, 1986), computational (Burton et al., 1990), and neuroanatomical (Gobbini and Haxby, 2007) models of person identification propose a sequential process, whereby the visual properties of faces are first analyzed to support recognition, followed by access to biographical details, such as social relationships, to aid identification. Models differ in the way that knowledge of the social relationships of an individual are stored and accessed. According to one proposal, social relationships are stored as general semantic memory, and accessed by the activation of a person-identity node (Burton et al., 1990). In contrast, others have suggested that information about the associations between people are directly linked to the representation of each individual (Barry et al., 1998; Wiese and Schweinberger, 2011). Priming presentations of social connections by viewing the face of an associate immediately before viewing a face modulates the N400 event-related potential, indicating that knowledge of social connections can be accessed at least within 300-400 ms (Wiese and Schweinberger, 2008, 2011). These priming effects are distinct from priming due to semantic relatedness (e.g., having the same occupation), and suggest a representation of connections between people we know that is accessible in the first few hundred milliseconds of observing them. Determining how these social connections are represented, how the neural basis of the process unfolds over time, and how it interacts with visual processing and the processing of individual identity, is an important step in furthering our knowledge of the neural underpinnings of person identification and social cognition.

To examine the temporal dynamics of neural activity associated with representing social connections, we used electroencephalography (EEG) and temporal representational similarity analysis (RSA) as participants viewed the faces of individuals from a social network. Participants first become familiar with the individuals and learned relationships between them in a naturalistic setting by viewing episodes of a television show in which the members interacted. We hypothesized that patterns of neural activity, evoked by viewing the faces of members of the network, would represent the connections between each member. Our data also allowed us to reveal how the representations of social connections evolve over time, and their relationship to the visual processing of each face.

## Materials and Methods

### Participants

33 right-handed adults with normal or corrected vision (10 males; age range 18-29; average age = 21; ethnicity = 47% Asian, 5% Black/African American, 16% Hispanic/Latinx, 26% White/Caucasian, 5% Other) participated in this study. They were recruited from the George Mason University psychology research participation pool, flyers posted on campus, emails, and social media posts. Participants signed a consent form in accordance with the Declaration of Helsinki and the Human Subjects Review Board at George Mason University and were compensated for their time through money or course credit.

### Task Design

#### Behavioral Task

Participants watched three episodes of NBC’s Parks and Recreation (63 minutes total; 21 minutes each) through the video streaming service Netflix. All subjects reported that they were naïve viewers of the episodes on a pre-screening survey prior to participation. Through viewing, they became familiar with a social network consisting of 10 characters. The characters in the network are friends and coworkers with varying levels of social closeness, as shown through the quantity and quality of social interaction throughout the episodes.

This method of social network exposure was chosen to be realistic to how social relationships are observed in the real world in a variety of situations. Previous studies in the authors’ lab used a paired association learning task to familiarize participants with social networks, with frequency of face image pairing indicating closer relationships (Dziura and Thompson, 2019). Frequency of association is highly related to social closeness, but it is not necessarily the same thing (Tarr et al., 2016), so this more naturalistic observational method was chosen to incorporate interaction quality as well as quantity.

After watching the show, participants filled out a survey where they reported several relationship characteristics among the network. These included: 1) how frequently each character interacted, 2) how much they like each other, 3) how similar their personalities are, and 4) based upon all other factors, how close they are to each other. This final question was used to create models of perceived social closeness to include in the analysis. The questions were asked on a scale of 0-6 (except for the liking question, which ranged from -6 to 6) where higher numbers indicate higher/greater relationship characteristic. Interaction time for each character pair was also collected by counting the time (in seconds) that each character was in a scene with the other character. This included group interactions, not only the time characters directly spoke to each other one-on-one (e.g. some scenes had one character talking to several others, or a number of people all talking in sequence). The interaction times of character pairs ranged from 0 (they were never actually seen to interact through the three episodes) to 450 seconds (see **Figure 2F** and **Figure 3A**).

**Figure 1.**
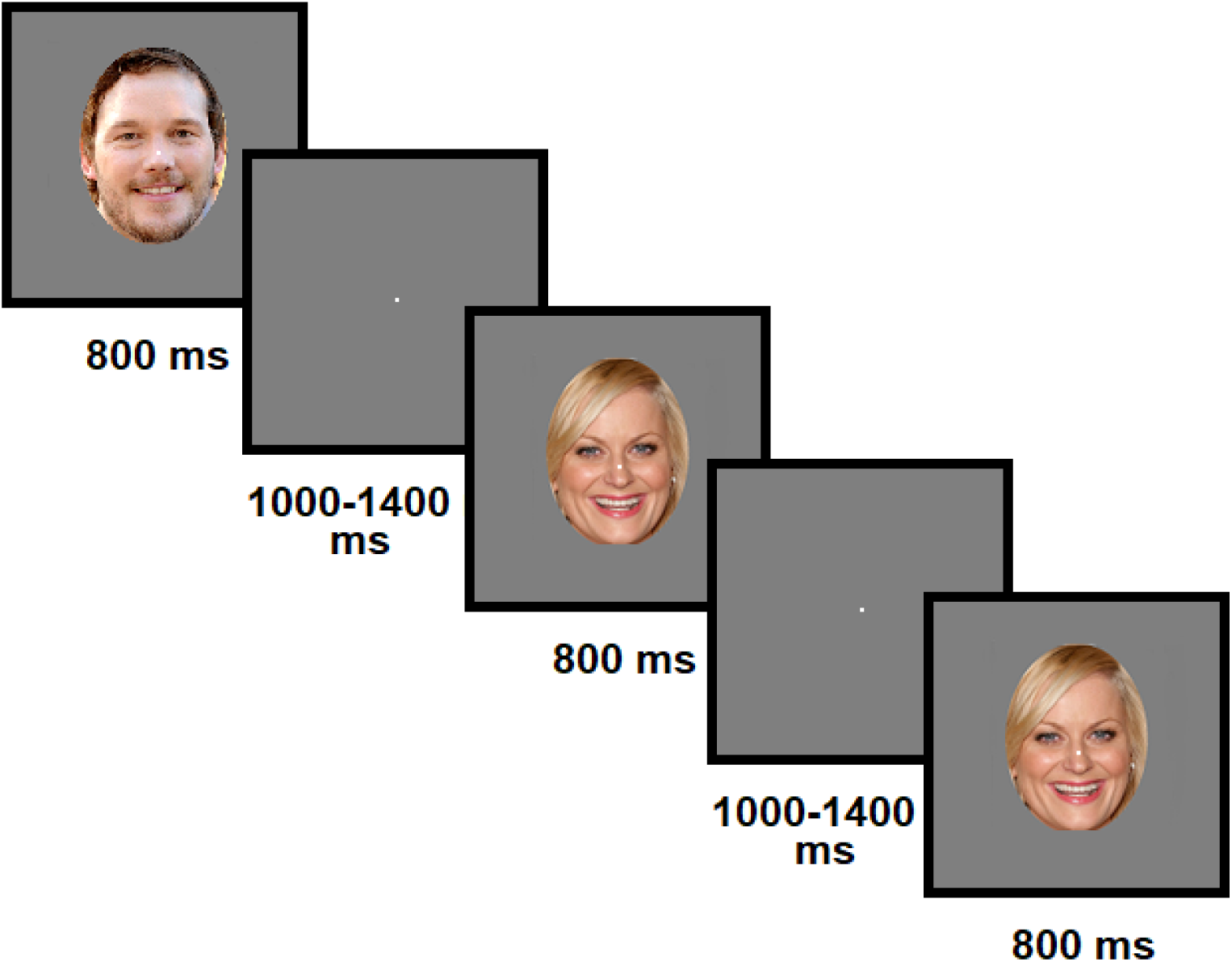
Experimental design. Faces were presented centrally one at a time while participants engaged in a 1-back task.

**Figure 2.**
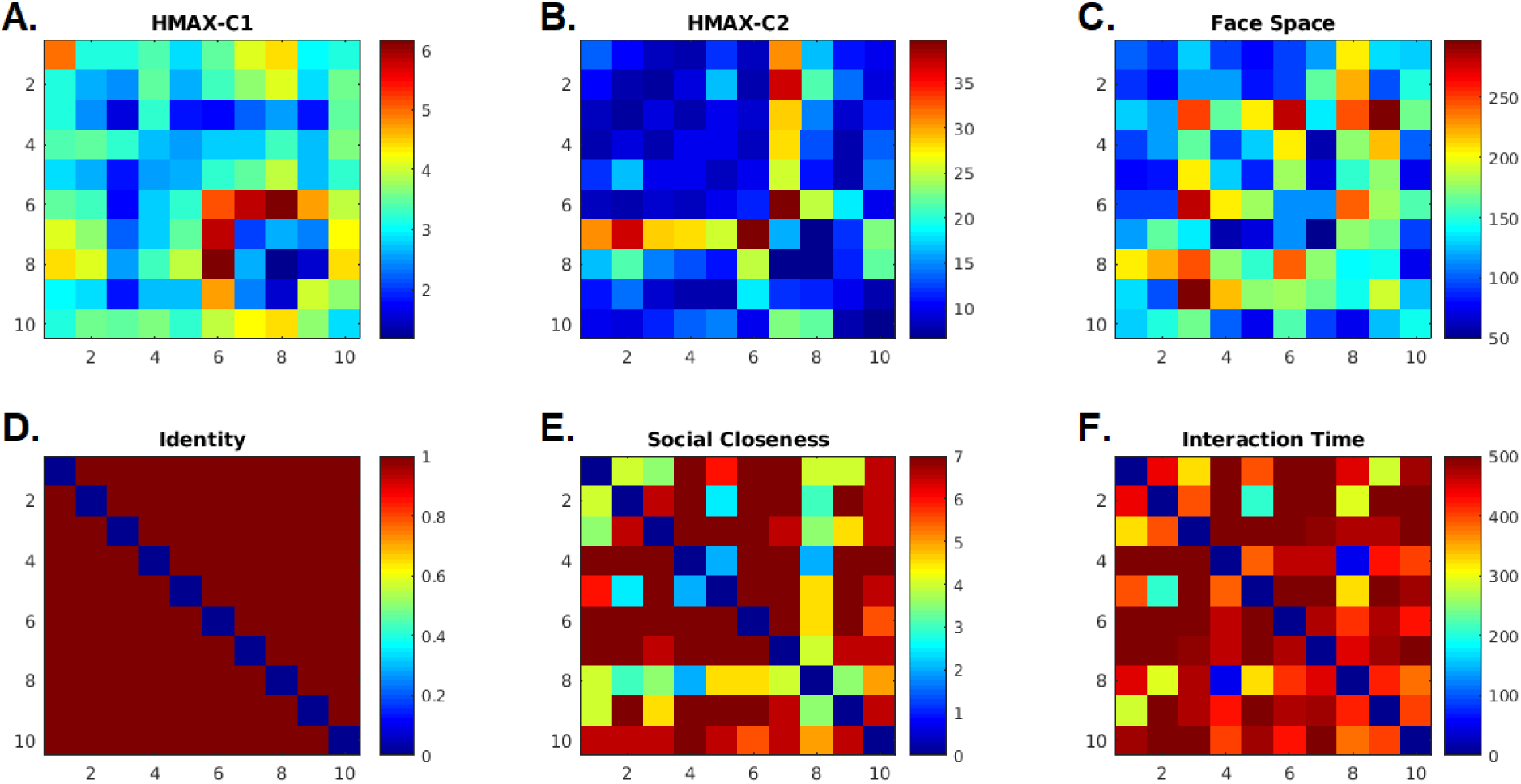
Dissimilarity matrices across a change in facial expression for each character. **A-B.** Euclidean distance from the C-1 and C2 layer outputs of the HMAX model of visual cortex responses. **C.** Euclidean distance in representational face space. **D.** Identity of the characters, with 0 on the diagonal indicating different images of the same characters and all other pairings as equally dissimilar. **E.** Social distance converted from a sample subject’s survey responses. **F.** Interaction time (inverted as 700 – true time) in seconds between each character.

**Figure 3.**
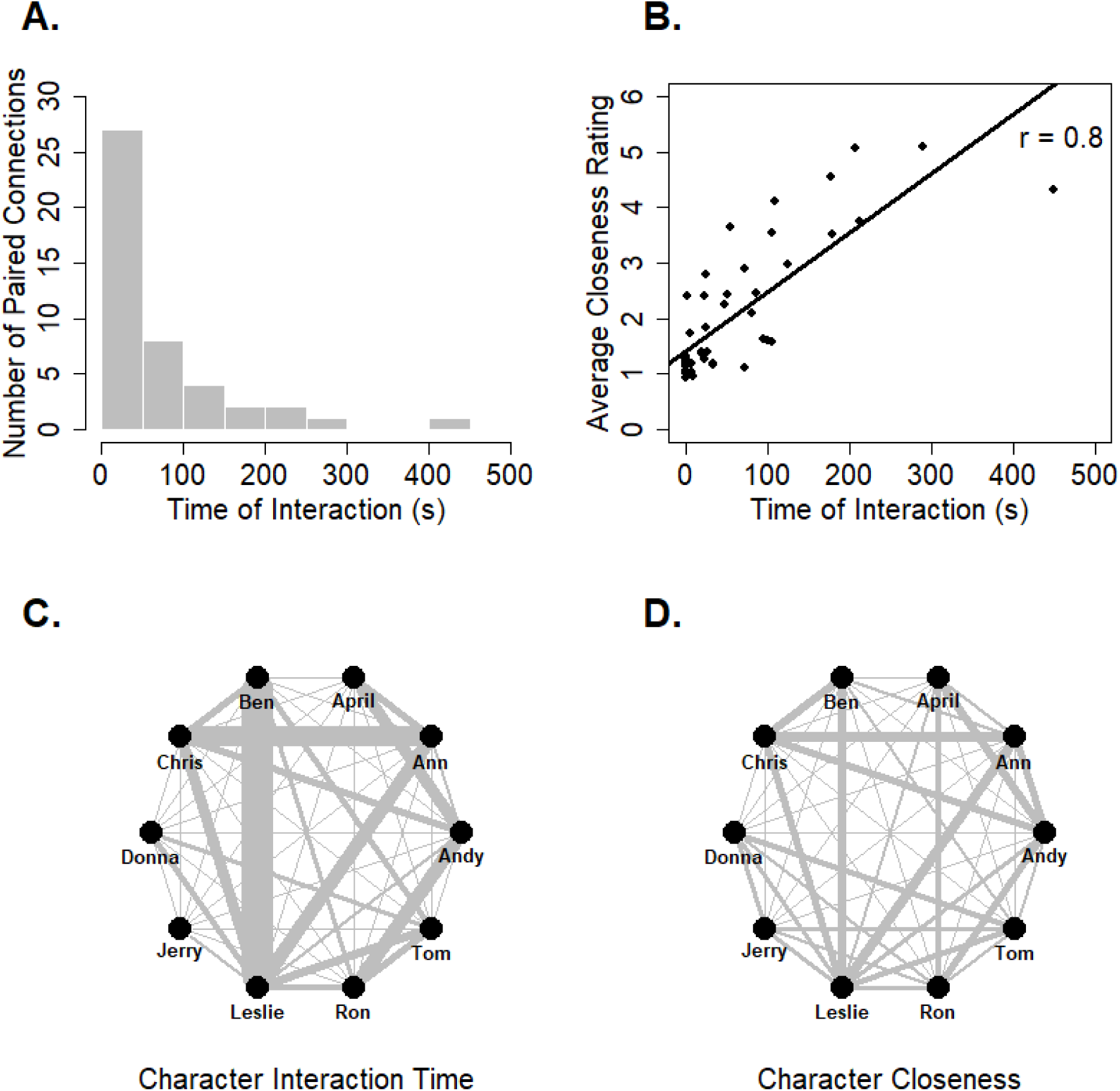
Specifics of the 10-person social network observed by participants. **A.** Histogram showing the distribution of times the characters spent interacting. **B.** Correlation between the time characters spent interacting and the averaged social closeness rating. **C.** Network map using interaction time as connection weight. **D.** Network map using the averaged social closeness rating as connection weight.

#### EEG Task

After the behavioral tasks, participants completed an EEG session where they viewed images of each of the 10 characters. Stimuli consisted of 2 color images of every character, each presented a total of 96 times across the entire session. The two images of each character differed in facial expression, and all images were collected from an internet image search (rather than by taking screenshots of the episodes). Each image was presented for 800 ms with a variable inter-stimulus interval ranging from 1000-1400 ms. Participants completed a 1-back task, where they pressed a button after an immediate repeat of an image (**Figure 1**). This was to ensure visual attention, and neural data from the repeated images were not included in further analysis. The screen background was grey (RGB = 127) with a constant white fixation dot. 8000 ms rest periods were presented after every 21 images, and a self-paced break period was presented after 2 blocks to ensure that participants did not get overly fatigued and had time to rest their eyes.

### Data Acquisition and Analysis

#### EEG Acquisition

EEG signal was recorded with 64 active Ag/AgCl electrodes (Brain Products actiCHamp system; http://www.brainproducts.com/) placed on the head according to the modified 10-20 system (American Electroencaphalographic Society, 1994). Four extra channels were placed around the eyes (two near the outer canthi of each eye and two above and below the left eye) to monitor horizontal saccades and vertical eyeblinks. A final electrode was placed on the nose to use as a reference. All EEG electrode impedances were reduced to <50 kΩ prior to beginning the experiment, and the signals were low- and high-pass filtered online between 0.1 Hz and 249 Hz and digitized at 500 Hz.

#### Data Pre-processing

All EEG pre-processing was completed in Matlab R2017a (Math Works, Natick, MA; https://www.mathworks.com/). Initial steps were conducted using the EEGLAB and ERPLAB toolboxes (Lopez-Calderon and Luck, 2014; Delorme and Makeig, 2004). Signals from electrodes with a high amount of noise were removed and replaced with interpolated data from surrounding electrodes. Data was bandpass filtered between 0.5 and 30 Hz using a non-causal Butterworth impulse response function to remove very low and very high frequency noise. A combination of artifact correction (using independent components analysis) and artifact rejection was conducted. Components that were a) indicative of noise and b) correlated with EOG channels (indicative of blinks) were selected for rejection using SASICA (Chaumon et al., 2015) and subsequently subtracted from the data. Following this, any trials remaining with EEG signal +/-100µV in any channel were removed from the data (Sawaki et al., 2015).

Data was separated into epochs ranging from -200 ms pre-stimulus onset to 600 ms post-stimulus onset and averaged within conditions (each individual face image and 1-back repeats). This yielded a single datapoint for every electrode for every timepoint (2 ms apart) from -200 to 600 ms for each condition. Pre-stimulus onset data was checked to ensure that there were no significantly >0 correlations.

#### Representational Similarity Analysis

Representational similarity analysis (RSA) is used to compare between-stimulus similarity across modalities that may be very different on the surface (Kriegeskorte et al., 2008). In this case, averaged ERPs from 64 scalp sites are correlated with proposed models of different types of information suspected to be involved in face perception. Six dissimilarity matrix (DMs) models representing three types of information were used to this end: visual properties, character identity, and social relationships (**Figure 2**).

All models were compared to neural distance across two different facial expressions within each character identity (Vida et al., 2016). The primary model of visual properties was the Euclidean distance between responses of the C2 layer of HMAX, which simulates the complex visual cell response to an input image (Riesenhuber and Poggio, 1999). Character identity was modeled as either the same (0) or different (1) across the two images of each character (Vida et al., 2016). The social relationship between each character was measured using subject responses to a questionnaire administered after the video viewing portion (“social closeness”). Three other models were tested to strengthen the evidence for each category of information. Additional visual models included the Euclidean distance between responses of the C1 layer of the HMAX model and the distance of faces in face shape PCA space (Kramer et al., 2016). This latter model was created using Interface (Kramer et al., 2016) by placing 82 fiducial landmarks on each face. PCA was then performed on the set of faces, and the distance between each face in PCA space was calculated. The DMs created from the two layers of the HMAX model are correlated but not identical (Pearson’s r = 0.46, p < 0.001), while the DM created from the face space model is uncorrelated with either HMAX layer (r = -0.11, p = 0.39 / r = -0.15, p = 0.27). Finally, a social relationship model based on interaction time between each character was created. Unlike the social closeness model (which was created from subject-specific questionnaire responses), this was identical across all subjects because they all watched the same episodes.

#### Statistical Analysis

Null hypothesis testing was conducted through permutations at both the single subject and group level as per the recommendation of Stelzer et al. (2013). 100 random permutations of observation order (without replacement) were conducted for each correlation test for each subject as an intermediate step. Group average permutations were then conducted by randomly sampling one permuted dataset from each subject and averaging across all subjects. This was repeated 10,000 times (with replacement) to obtain a chance distribution map at each timepoint. Thresholds were assessed by finding the permuted datapoints on both ends of the distribution map that correspond to 0.1% chance (p = 0.001, two-tailed) for the separate models and 5% chance (p = 0.05, two-tailed) for the partial correlation analysis. The datapoints from the true similarity analyses that fall beyond those threshold points are considered significantly different from chance. Statistical analysis was performed in Matlab R2017a and timecourse figures were created using R version 3.5.1 (R Core Team, 2018).

## Results

### Survey Results

The social closeness questionnaire yielded pairwise measures of perceived interaction frequency, liking, personality similarity, and social closeness. All measures were significantly correlated with each other, as was an average of the first three questions (an implied measure of social closeness) and the explicit social closeness question (**Table 1**).

**Table 1.** Pearson correlations among the measures in the perceived social network survey. The bottom row in particular shows the correlations between the measure of social closeness ultimately used for further analysis and the other survey items, as well as an average of three items used previously to elicit social closeness (Tarr et al., 2016). All correlations are significant at *p* < 0.00001.

| Social Network Survey Measures |  |  |  |  |
| --- | --- | --- | --- | --- |
|  | Frequency | Liking | Personality Similarity | Average of Frequency, Liking, and Similarity |
| Liking | 0.79 |  |  |  |
| Similarity | 0.80 | 0.81 |  |  |
| Social Closeness | 0.91 | 0.85 | 0.77 | 0.92 |

Through these measures, two weighted networks were created with varying degrees of closeness among members (**Figure 3**). Every character was connected to every other character, as they are all presumed to know each other. However, within the three episodes, some characters interact much more often than others (**Figure 3A**), and relationships are therefore shown to develop more strongly among some members compared to others. The time in seconds that each character spent interacting across the three episodes ranged from 0-450 and yielded a network (**Figure 3C**) with density of 63.2 and eigenvector centralization index of 79%. The social closeness network (**Figure 3D**) ranged from 0.9-5.1 (scale of 0-6), with a density of 2.1 and eigenvector centralization index of 26%. These two network measures were highly correlated (r = 0.8, p < 0.001; **Figure 3B**).

### Event Related Potentials

As all EEG data was collected in response to face stimuli, we did not conduct any significance tests relative to a non-face control. However, the N170 face effect has been well documented (Bentin et al., 1996; Rossion, 2014) and we therefore expected to see large negative deflections around 170 ms for our stimuli. We were also particularly interested in the time around 100 ms, important for visual perception and potentially sensitive to faces (Colombatto and McCarthy, 2017; Dering et al., 2011; Liu et al., 2002) and 250 ms, another time period linked to differences in face perception (Huang et al., 2017). **Figure 4** shows example ERP data averaged across the entire group. At a temporoparietal electrode where we might expect to see face-relevant activity, there are characteristic positive and negative deflections around 100 and 170 ms respectively (**4D**). These images also show minor differences in the three waveforms for each chosen face stimulus, which suggests that there are potential meaningful differences in the data related to each individual face. The subject-specific waveforms from this data were subsequently used in RSA models to examine the representational relationships among the faces.

**Figure 4.**
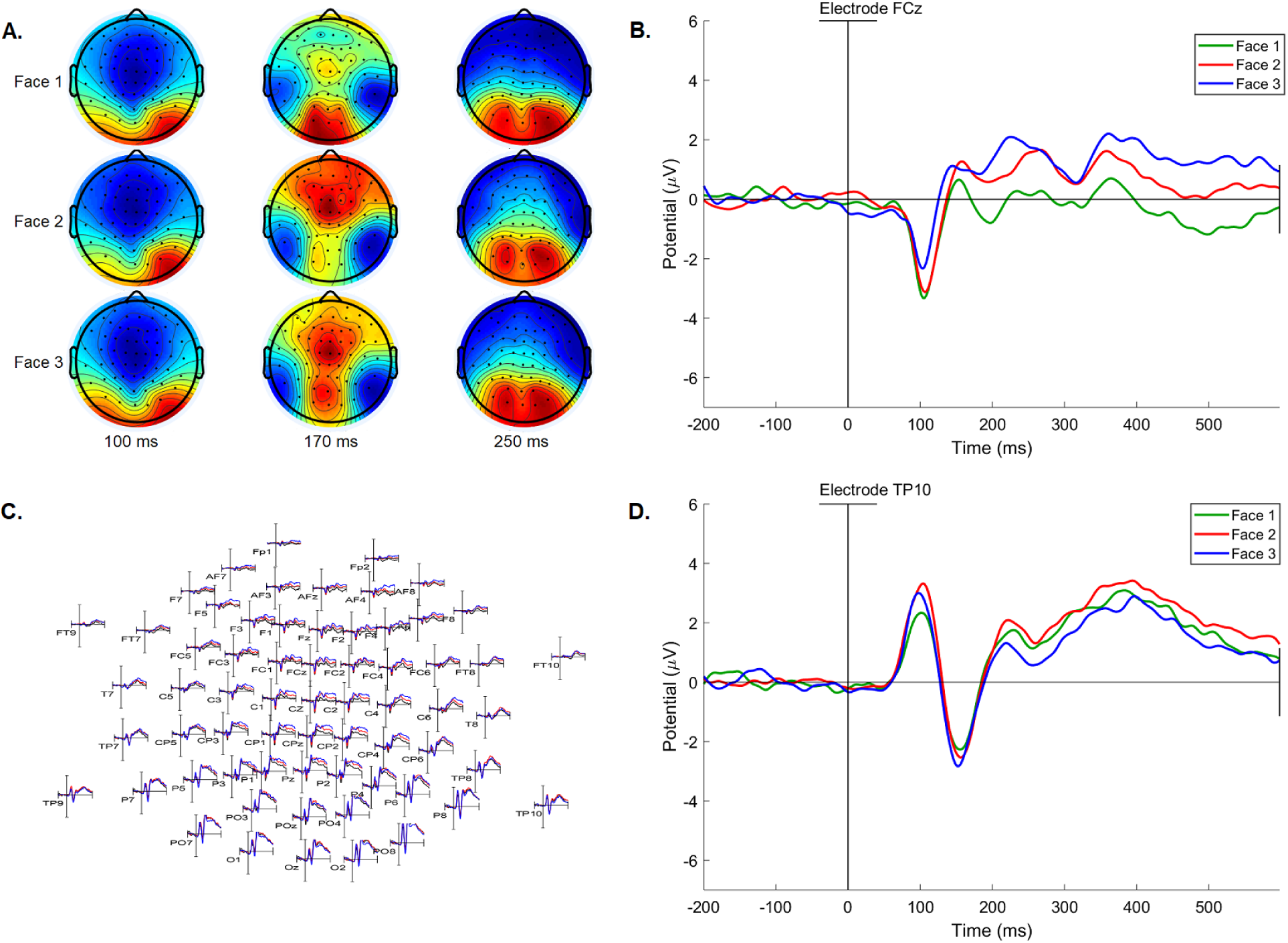
Example event-related potential data from three of the network face images averaged across all subjects. **A.** 64 channel scalp maps at three different times (100 ms, 170 ms, and 250 ms). **C.** Topographic map of every grand average ERP timecourse from -200 – 600 ms post image onset for the 64 electrodes. **B, D.** Larger waveform images from a fronto-central electrode (**B**) and a right temporoparietal electrode (**D**).

### RSA Results

All model types showed significant associations with neural patterns at different time points after image onset (p < 0.001) (**Figure 5**). The significant associations showed the expected pattern, with visual information becoming significant first (**Figure 5A**: C1: 96-232 ms / C2: 86-196 ms / face space: 68-238 ms), followed by non-visual social information (**Figure 5C**: closeness: 232-298 ms / interaction time: 210-304 ms). The cross-expression identity model was significantly associated with neural patterns both early and later, between 60-380 ms (**Figure 5B**).

**Figure 5.**
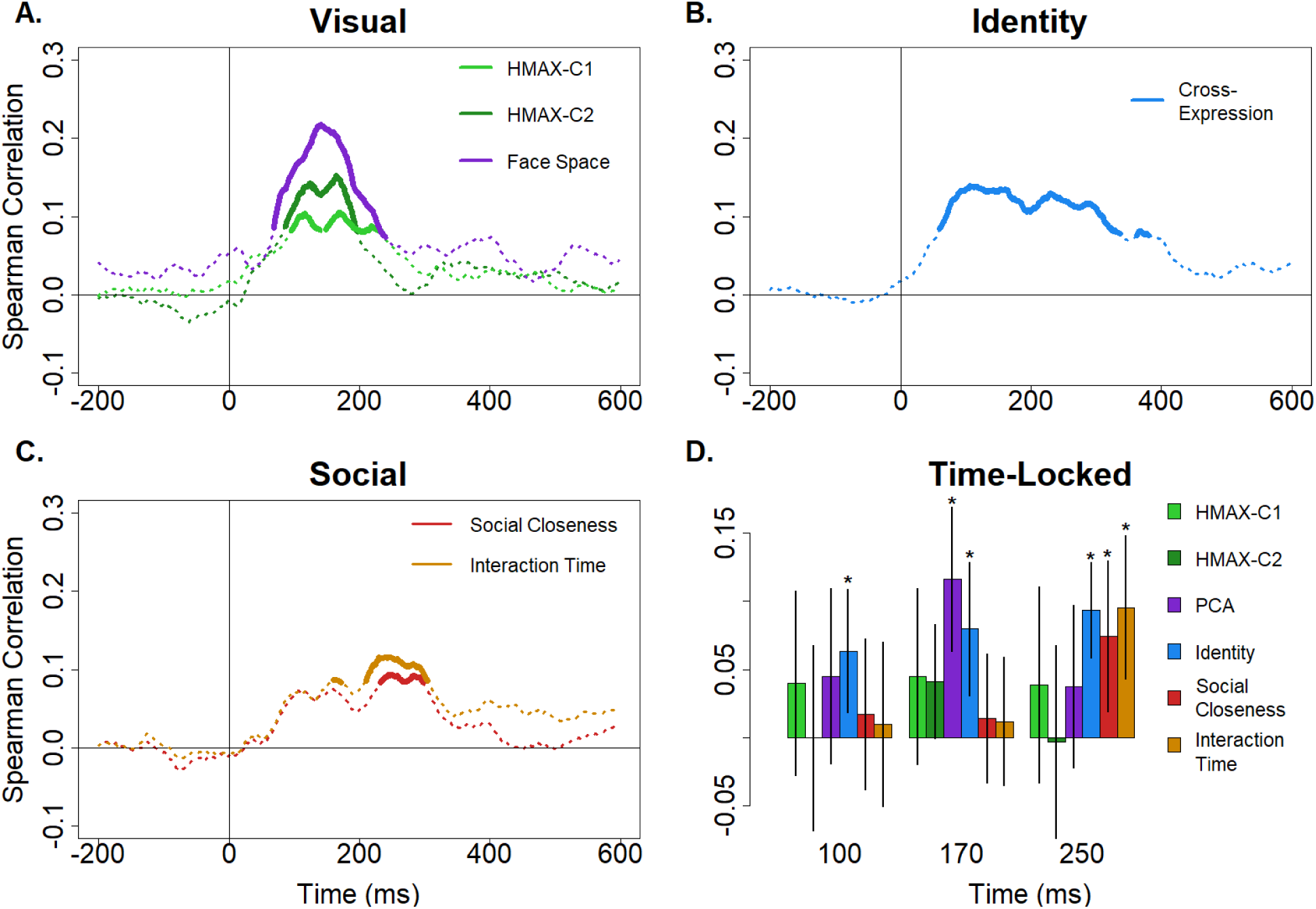
**A-C.** Full timecourse of each model’s correlation with neural pattern similarity from - 200 to 600 ms post-stimulus onset, organized by information type. Bold line sections are significantly greater than zero at p < 0001. **A.** Visual models, including two levels of HMAX and face space distance. **B.** Identity across a change in facial expression. **C.** Social models, including social closeness as measured by the social network survey, and time of interaction in seconds across the three episodes. **D.** Correlations between each model and neural pattern similarity time-locked to three time points associated with face processing (100 ms, 170 ms, and 250 ms). Significance and error bars reflect 95% confidence intervals.

To directly assess whether these models reflect information associated with typical ERP face-sensitive components, we also conducted time-locked analyses at 100 ms, 170 ms, and 250 ms. For each electrode, the data from each stimulus was averaged over 40 ms (20 pre- and 20 post) to get a static whole-head neural pattern for each face. These patterns were then correlated with the same DMs used in the previous analysis. Results are shown in **Figure 5D**. Neither HMAX layer was significant at 100 ms, which could reflect inter-subject variability in timing around early visual components (Guthkelch et al., 1987; Verma and Kooi, 1984), especially as both were shown to be significant with the full sliding window analysis. The face space model was significant at 170 ms but neither other time, and both interaction time and social closeness models were significant at 250 ms. As suggested from the original analysis, the identity model was significant at all 3 timepoints.

Partial correlation analyses were also conducted with DMs for visual, identity, and social connection. We found that all three types contribute unique variance to neural signals after a face is seen, although at a lower significance threshold (p < 0.05) (**Figure 6**). As with the separate model correlations, neural representations move from visual early to social later. However, a difference between the subject-specific model of social closeness and the generalized interaction time model emerged. When only visual and social inputs were included, both models contributed unique significant information to neural patterns. However, when the identity model was included, the social closeness model became non-significant. This pattern was not observed when interaction time was used as a measure of social closeness. In that instance, the significant time period for representations of interactions among characters was between 412-450 ms. In that analysis, the visual (HMAX-C2) model was significant from 76-202 ms, and identity was significant both from 34-384 ms.

**Figure 6.**
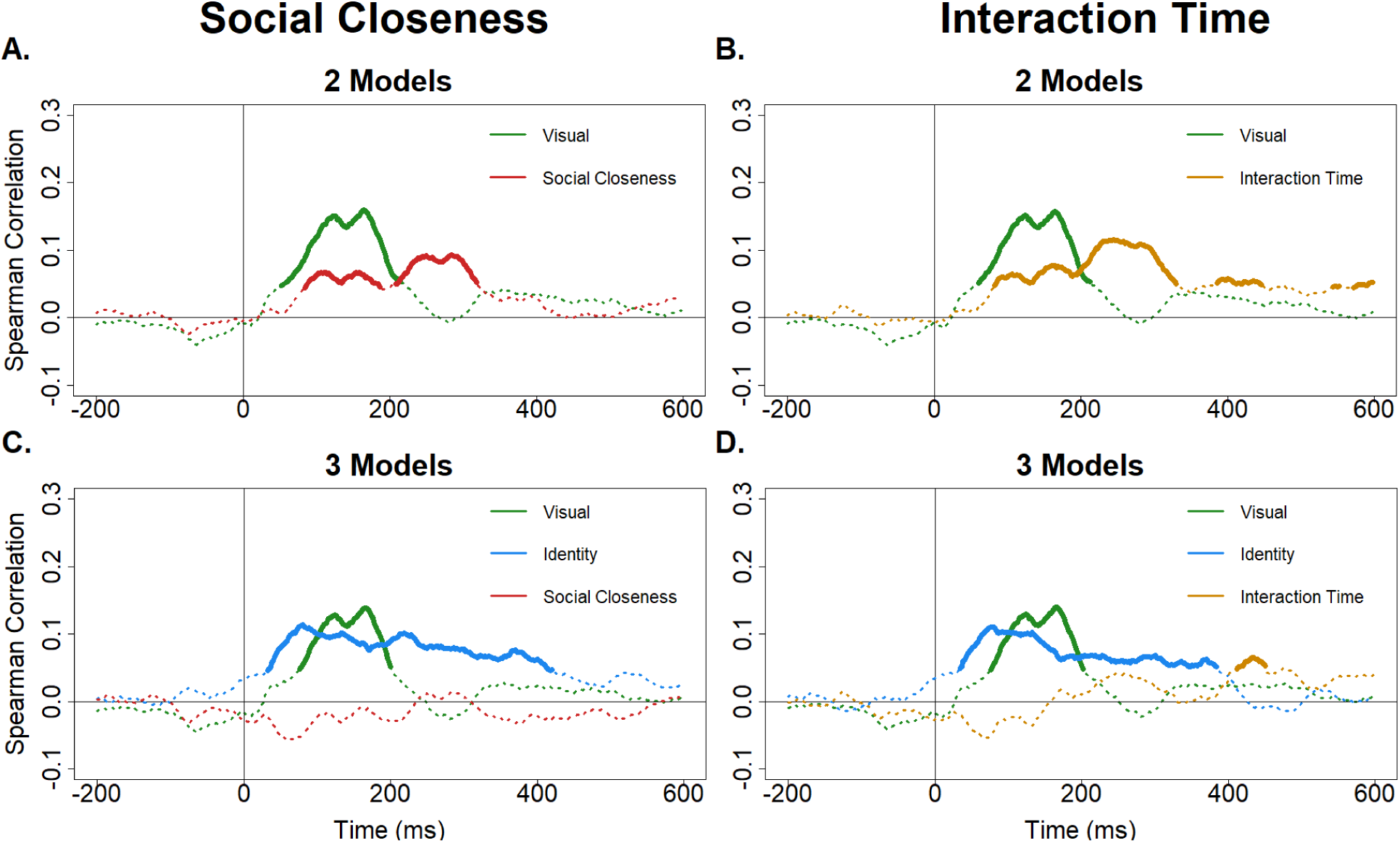
Spearman partial correlation analyses to examine the unique contributions of different types of information. **Top row**: Visual and social inputs. **A.** Partial correlations between the HMAX-C2 and social closeness models with neural pattern similarity. **B.** Partial correlations between the HMAX-C2 and interaction time models with neural pattern similarity. **Bottom row**: Visual, identity, and social inputs. **C.** Partial correlations between the HMAX-C2, identity, and social closeness models with neural pattern similarity. **D.** Partial correlations between the HMAX-C2, identity, and interaction time models with neural pattern similarity. Bold line sections are significantly greater than zero at p < 0.05.

## Discussion

This study examined the neural timecourse of the representation of familiar faces who were part of the same social network. Consistent with previous research, visual and identity information were present in ERP patterns across the whole brain early after image presentation. We also found that information reflecting social relationships is present in these whole-brain patterns. Our data indicate that progression of information content moves from simpler visual properties to more complex non-visual social information, and both of these are present at the same time as representations of identity. Furthermore, each type of information contributes uniquely to the neural patterns elicited by these faces at different points in the processing stream.

Different types of visual information are represented in the first 100 ms or so after face images are presented. However, by around 200 ms different models of visual processing of faces are no longer coupled with neural patterns. This is true for both layers of the HMAX visual cortex computational model and a model of the faces in principal component analysis face space, even though the HMAX and face space models are theoretically distinct, and not correlated with each other. The differences between the HMAX and face space models might reflect the more local nature of the former representation, whereas the PCA space model reflects the contribution of shape features, including global shape. The early neural representation of the visual similarity of faces is consistent with prior literature showing MEG and EEG responses to face stimuli at this time (Seeck et al., 1997; Schendan et al., 1998; Liu et al., 2002). A model of character identity across a change in character image is also clearly represented in neural patterns, which has been previously demonstrated using MEG and a similar cross-identity expression model (Vida et al., 2016). This model was associated with neural patterns for the longest period of time out of all the models, indicating that representation of person identity might encompass a number of underlying processes. Information about perceived social connections between individuals in a social network is represented in the brain as early as 230 ms after a face is presented, and peaks around 250 ms. Previous literature has linked a negative deflection around this time (N250) with accuracy in recognizing faces (Huang et al., 2017). Differences in neural activity in response to ingroup vs. outgroup images have also been shown to peak around 250 ms (Ito et al., 2004). Priming of a face by known associates of an individual has been previously been reported to modulate the N400 ERP (Wiese and Schweinberger, 2008, 2011). Our data suggest that representations of the strength of connection between individuals who belong to the same social network are encoded as early as ∼250 ms after first viewing the face.

The results of our study show that face identity is not solely driven by low level visual properties, as captured by the HMAX model, or face shape as captured in PCA space. A recent study indicated that gender and age information is also encoded early in the MEG response to faces, especially familiar faces, and overlaps with representations of identity (Dobs et al., 2019). Other recent work has highlighted the role of surface texture on the neural representation of face identity (Nemrodov et al., 2018, 2019). Surface features and invariant shape properties associated with gender and age might be expected to contribute to (dis)similarity in HMAX and/or face space (Johnston et al., 1997), although clearly these models do not capture the entirety of early representations of face identity. Modeling identity and social connection information separately showed that both representations are present later in the neural response. When the model of identity was included with subjective ratings of the social closeness of individuals, the neural response explained by the latter was no longer significant in the 200-400 ms period.

Neural responses around this time have been related to explicit categorization of faces according to gender, familiarity, and group membership (Barrett and Rugg, 1989; Jemel et al., 2009; Olivares et al., 2003; Valdes-Conroy et al., 2014). Taken together, these results suggest information reflecting social connections that is not separable from person identity is present early, potentially contributing to recognition of faces, and that identity-independent information associated with social connections may be a separate process that occurs only after identity representations are processed.

The divergence between the social closeness and interaction time models could reflect inter-subject variability in the encoding of social relationships from the episodes that they watched, or else could reflect the difference between interaction quantity (time spent interacting among people) and quality (how the people actually feel about those interactions). The social network learning task was brief compared to observations of social interactions that people might generally have among groups in the real world. Research into personal social networks suggests that information about familiar people is stored or represented in the medial prefrontal and anterior temporal cortex (Krienen et al., 2010; Thornton and Mitchell, 2017; Wlodarski and Dunbar, 2016). There is likely a difference between personally relevant social connection information (such as the relationships among kin or close friends) and non-personal social connection information (such as these relationships among acquaintances or even friends of friends) above and beyond mere stimulus familiarity (Gobbini et al., 2004; Keyes and Zalicks, 2016; Krienen et al., 2010). Further research is needed to examine the differences between quantity time observed (familiarity) and potentially more meaningful qualities of a relationship in the context of larger interconnected social groups.

Importantly, these neural patterns are elicited even without explicit directions to think about the identities or social relationships of the characters. This strengthens evidence from previous studies that show spontaneous neural coding of network characteristics within participants’ own social networks (Parkinson et al., 2017). The social functional theory of face processing posits that attention is naturally allocated to the most relevant cues, which in the case of the human perceptual system, includes social information (Adams et al., 2017). This would suggest the possibility that our participants were either implicitly or explicitly thinking about these relationships without us directing them to do so.

The speed at which this spontaneous process happens has implications for a number of potential behaviors. The more quickly we understand the purpose of another’s actions, the better able we are to react immediately in uncertain or dangerous situations. Information acquired from faces can give near-instantaneous cues toward conspecific threats (Öhman, 1986). Negative emotions present in faces have been linked to increased early visual attention around 120 ms, similar in timing to neural responses to images of snakes, which are thought to be evolutionarily relevant (Langeslag and Van Strien, 2018). Social information such as trustworthiness is also automatically tracked when a face is seen even in the absence of an explicit judgment task, and these traits may be fully processed by 100 ms (Engell et al., 2007; Todorov et al., 2009; Willis and Todorov, 2006). The results from this study suggest not only that the social information gathered from faces includes representations of relationships, but that this process happens within the first half second of perception. Similar to knowledge (or perceptions) about trustworthiness, knowledge about social relationships may be able to aid in rapid judgment processes. For example, if we understand the connections between people in social groups, we can more easily trust or rely on information that is passed among certain group members.

A significant part of a person’s identity relates to the social connections they form among others in groups. People often self-select into different groups or use group categorizations to form their own identities (Amiot and Aubin, 2013), and our identity is in turn often shaped by the people we surround ourselves with (Guo and Li, 2016; Van Veelen et al., 2013; Vivona, 2013). How these in-group connections interact with each other and relate to the individual identities that make up the group, as well as how they are learned and represented by those outside the group, is important to understand for people living in a social world.

## Acknowledgments

We thank Craig McDonald and Martin Wiener for providing critical input when designing this project. We also thank Leo Chase, Colleen Flanagan, Clara Glagola, Michael Norton, and Claudia Torresi for data collection assistance.

